# Proactive and reactive construction of memory-based preferences

**DOI:** 10.1101/2023.12.10.570977

**Authors:** Jonathan Nicholas, Nathaniel D. Daw, Daphna Shohamy

## Abstract

We are often faced with decisions we have never encountered before, requiring us to infer possible outcomes before making a choice. Computational theories suggest that one way to make these types of decisions is by accessing and linking related experiences stored in memory. Past work has shown that such memory-based preference construction can occur at a number of different timepoints relative to the moment a decision is made. Some studies have found that memories are integrated at the time a decision is faced (reactively) while others found that memory integration happens earlier, when memories were initially encoded (proactively). Here we offer a resolution to this inconsistency, demonstrating that these two strategies tradeoff rationally as a function of the associative structure of memory. We use fMRI to decode patterns of brain responses unique to categories of images in memory and find that proactive memory access is more common and allows more efficient inference. However, we also find that participants use reactive access when choice options are linked to a larger number of memory associations. Together, these results indicate that the brain judiciously conducts proactive inference by accessing memories ahead of time when conditions make this strategy more favorable.

## Introduction

Some decisions are made repeatedly, offering the opportunity to learn directly about an option’s value through past experiences with its outcome. However, decisions often consist of a choice between options whose outcomes have not been directly experienced before. Computational theories of planning suggest that one way to approach such decisions is by knitting together separate relevant memories through mental simulation^1–3^. The ability to flexibly combine information in this way is central to intelligence: it frees us from having to decide based on direct trial-and-error experience alone and enables us to make inferences and to plan novel courses of action using cognitive maps or internal models^4–8^.

The process of drawing inferences requires accessing relevant memories and recombining or integrating across them to build new relationships. Studying memory access is therefore one way to shed light on the covert mechanisms that give rise to inferential choice. Yet previous work attempting to probe this connection has left open a critical gap in our understanding of how and when memory integration supports inference. In particular, some studies have claimed that memories are accessed at the time a choice is faced^2,9,10^, while other studies have found that memory access occurs much earlier, when relevant memories are first encoded^11,12^. These two approaches differ not just in the timepoint of memory access, but also reflect distinct mechanisms. Integrating memories during a decision requires “on the fly” processing, which is likely to take time, whereas integrating memories earlier suggests that the new model for inference already exists when a choice is later made, yielding more efficient decisions^11,13,14^. It has been suggested, but not yet empirically tested, that there may be some normative explanation for the variation between these two approaches^15^. In the present study, we aimed to address this gap by studying both possibilities in a single experimental design. We sought to first confirm the normative advantages that early memory access confers and then to investigate how changing the structure of memory access can rationally shift this process to happen later, at decision time.

The role of memory integration in inference is often studied with multi-phase tasks that first seed relevant associative memories and then test whether people integrate them when probed to make decisions. A classic task in this vein, which we build upon here, is *sensory preconditioning*^16^. In sensory preconditioning, participants are first trained to associate two stimuli that occur in succession (A→B). Then, in a separate phase, the B stimulus is associated with reward. The critical question is whether people infer that the A stimulus is also associated with reward. This is tested in the final decision phase, when participants are asked to choose between A and another control stimulus (which is equally familiar but lacks the indirect reward association). Humans and non-human animals alike tend to prefer A despite never directly experiencing its association with reward^11,12,14,16^. Studies of sensory preconditioning and similar tasks have revealed two potential mechanisms, each predicting memory integration either before or during choice, that may lead to this same behavioral effect.

A typical explanation for inference in tasks like sensory preconditioning, assumed in theories of decision making that date back to Tolman^8^, envisions that choosing A reflects prospective mental simulation at decision time: in this case, retrieving the B-reward association when evaluating whether to choose A. This, in turn, is thought to be a minimal case of a more general capacity for forward planning. This forward planning has been embodied by theories of model-based reinforcement learning in which actions are evaluated over multiple steps using a learned internal model, either in the form of one-step associations between states encountered serially or as a successor representation that generalizes this to associations over multiple timesteps^17–19^. By examining neural signatures of memory retrieval, it has been possible to investigate how memory access actually relates to successful model-based inference. Yet, studies have yielded mixed support for this account. Some evidence suggests that both humans and non-human animals engage in prospective retrieval at decision time, and that this pattern is associated with inferential performance^4,9,10,20–22^. However, there is also evidence that associative recall may occur long before a decision is ever faced^11,12,23–26^.

These latter findings imply a second explanation for inference in these tasks: that the value of options may be pre-computed when relevant information like reward is first encoded, thereby preempting the need for evaluating potential outcomes later at choice time. In some studies of sensory preconditioning, for instance, it has been found that when B is presented during reward learning, A is concurrently retrieved and directly associated with reward^11,12^. Such a strategy is feasible because, at this time, participants have already been provided with all of the components necessary to form a complete model of the task. Perhaps analogously, in rodent spatial navigation tasks, hippocampal place cells often briefly represent trajectories in front of the animal^20–22^, a potential substrate for prospective evaluation. However, otherwise similar “replay” events can instead reflect backward or altogether nonlocal trajectories at the time of reward^27–30^, potentially supporting a spatial analogue of the alternative inference strategy.

An emerging idea is that these different inference mechanisms may be special cases of a more general set of computations that share the common goal of integrating memories to infer action values, but that access memories at different times: either *proactively* before they are needed or *reactively*, once required for choice^15,31^. This in turn raises questions about how these strategies are balanced or adaptively deployed, and whether such control might explain variable results across studies. Indeed, the possibility of proactive computation implies that the brain must somehow be judicious about which memories it accesses, and when, since there are so many possible later actions that might be contemplated.

This idea, while compelling, is still largely untested, and raises a number of questions about how and when different strategies are deployed, which we aimed to address in this study. First, is it indeed the case that a proactive memory access strategy can support inferential choice equivalent to a reactive one? Second, what are the tradeoffs of the two approaches: if access occurs proactively, does it reduce the need for computation at decision time? Finally, do people rely differentially on this strategy at times when it would be sensible to do so?

We aimed to answer these questions by attempting to alter participants’ reliance on proactive inference. We had three primary hypotheses. First, we expected to confirm earlier (but inconsistently reported) results that sensory preconditioning can be solved with proactive memory access at the time of reward learning. Second, because proactive inference offers the advantage of a pre-computed value association, we hypothesized that this approach may allow for more efficient future decisions–i.e. decisions that are faster and more accurate. Third, we hypothesized that reliance on this strategy would adapt under different circumstances, which we operationalized by manipulating how difficult it is to access and integrate relevant memories. Drawing upon a rich tradition of research on associative memory^32^, we reasoned that having multiple relevant associations with an experience should, at any timepoint, induce competition between them, making their retrieval for use in inference less likely.

To test these hypotheses, we developed a novel learning and decision making task based on sensory preconditioning, and measured memory retrieval at multiple timepoints of this task while scanning participants with fMRI. Participants completed this task in three phases (**Figure 1**). In phase one, *stimulus learning*, participants learned associations between several antecedent-consequent (A→B) pairs of images. In phase two, *reward learning*, participants learned that a subset of consequent (B) images led to a reward, while others did not. Finally, in phase three, the *decision phase*, participants made a series of *test* and *transfer* choices between two of these images. On test choices, participants chose between consequent images that were directly associated with either a reward or neutral outcome during the reward learning phase. Transfer decisions consisted of choosing between antecedent (A) images that were paired with consequent images during the initial stimulus learning phase. Successful transfer of value to these images involves relying on memory for the paired association and can be accomplished, in principle, by either proactive or reactive memory access. This task is well suited to address our questions, which focus on when associations between memories are accessed to support inference. However, it is agnostic as to questions about how these associations are represented as internal models in the brain ( i.e. whether they are stored as one-step relationships or as a successor representation^17–19^).

**Figure 1.**
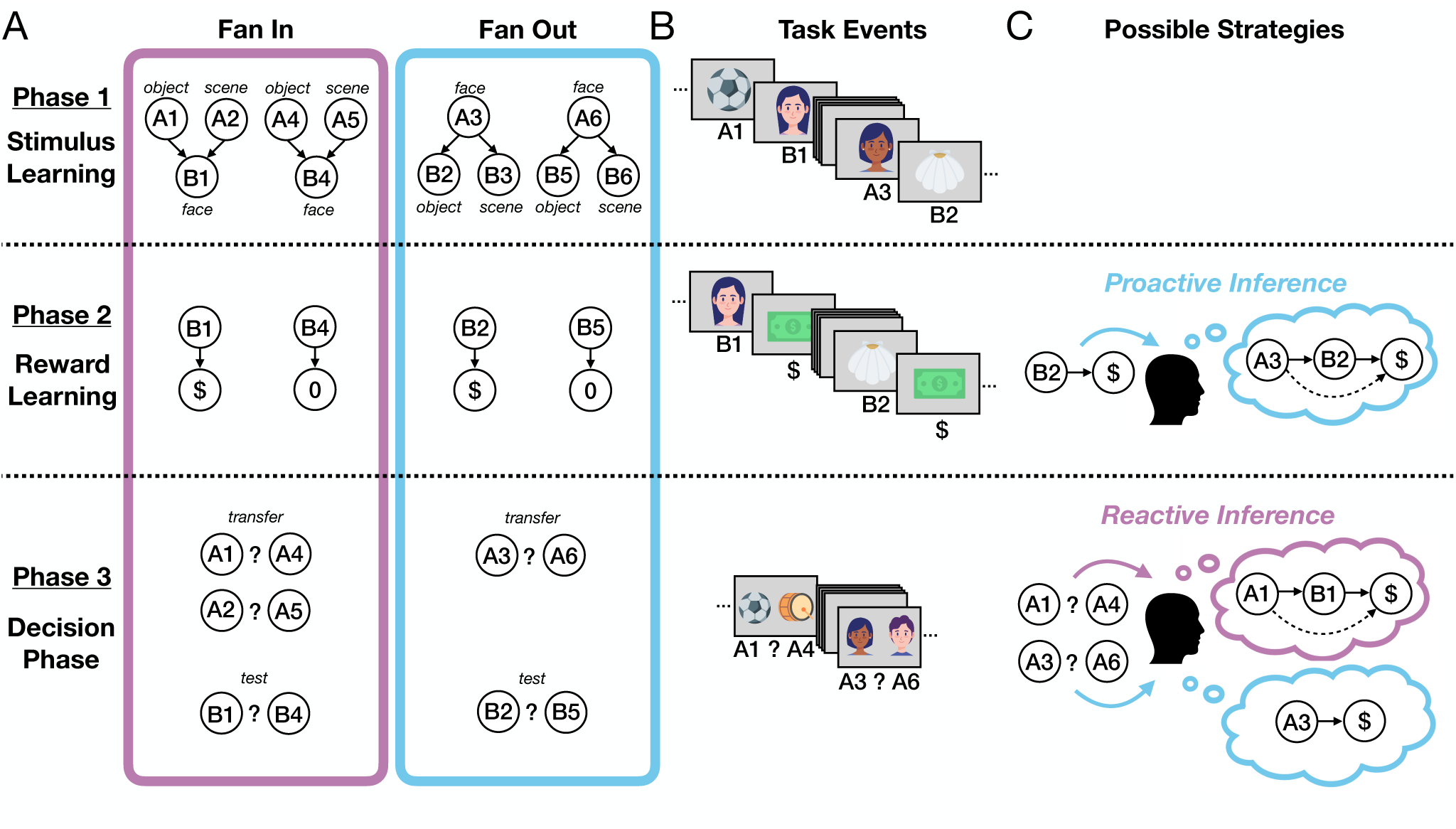
Task design and inference strategies. A) Task structure. Participants (n=39) underwent fMRI scanning while completing a three-part experiment with two different conditions, based on sensory preconditioning. The phases were similar for both conditions, which differed only in their specific associative structure. In phase one, *stimulus learning*, participants learned associations between several pairs of images (faces, scenes, or objects). Unknown to participants, there were two types of trials governing how these associations appeared. *Fan In* trials consisted of one of two possible antecedent A images followed by one consequent B image. *Fan Out* trials consisted of one antecedent A image followed by one of two possible consequent B images. Example categories for each image are shown here, and this was counterbalanced across participants. In phase two, *reward learning*, participants learned that a subset of consequent B images led to a reward, while others did not lead to reward. Finally, in phase three, the *decision phase*, participants chose between two images. Choices between consequent B images were used as *test* trials, whereas choices between antecedent A images were used as *transfer* trials. **B) Example events.** An example of the sequence of task events seen by participants in each phase. **C) Possible inference strategies.** Participants can engage in either of two inference strategies: proactive inference, at the time of reward learning, or reactive inference, at the time of the decision. During decision making, proactive inference does not require the integration of a memory with value, as this association has already been performed during reward learning. Due to differences in the number of competing antecedent memories at reward learning, we expected reactive inference to be used more for Fan In stimuli.

To capture putative reactivation of associations in memory in the service of inference, we exploited the fact that viewing different visual categories (e.g. faces, scenes, and objects) elicits unique activity in visual cortex^10,11,33,34^. We used images from these different categories for each of the different stimuli, which allowed us to measure whether reactivation of associated images in memory occurred during either reward learning, signifying proactive inference, or during decision making, signifying reactive inference. We predicted that proactive memory access during reward learning should result in more efficient later choices, and that reactive memory access during choice itself should have the opposite effect.

To address our third hypothesis specifically, we further varied the number of competing associations with a given stimulus by training participants on antecedent-consequent relationships under two different conditions (**Figure 1**). In one condition, two antecedent stimuli each predicted a single consequent stimulus; we refer to this as the *Fan In* condition. By contrast, in the *Fan Out* condition, a single antecedent predicted two possible consequents. The logic of this manipulation is that the Fan In condition induces greater retrieval competition between memories of antecedent stimuli when the consequent stimulus is presented during the reward learning phase. We therefore predicted that there should be increased reliance on reactive inference for stimuli in the Fan In condition relative to Fan Out condition. To test this prediction, we measured reactivation in BOLD activity for antecedent stimuli in the Fan Out condition during the reward learning phase, and for consequent stimuli in both conditions during the decision phase.

## Results

### Behavioral evidence for proactive inference and its modulation by retrieval competition

We first examined whether participants learned to directly associate consequent stimuli with reward, and whether they transferred value to associated antecedent images. To assess this, we analyzed participants’ test and transfer choices during the decision phase. On test choices, participants were highly accurate and tended to choose the rewarded consequent image over the neutral consequent image (β_0_ = 5.009, 95% *CI* = [4.085, 6.279]; **Figure 2A**). There was no difference between the Fan In and Fan Out conditions (β*_conditions_* = 0.321, 95% *CI* = [−1.251, 2.128]), indicating that participants learned similarly in both.

**Figure 2.**
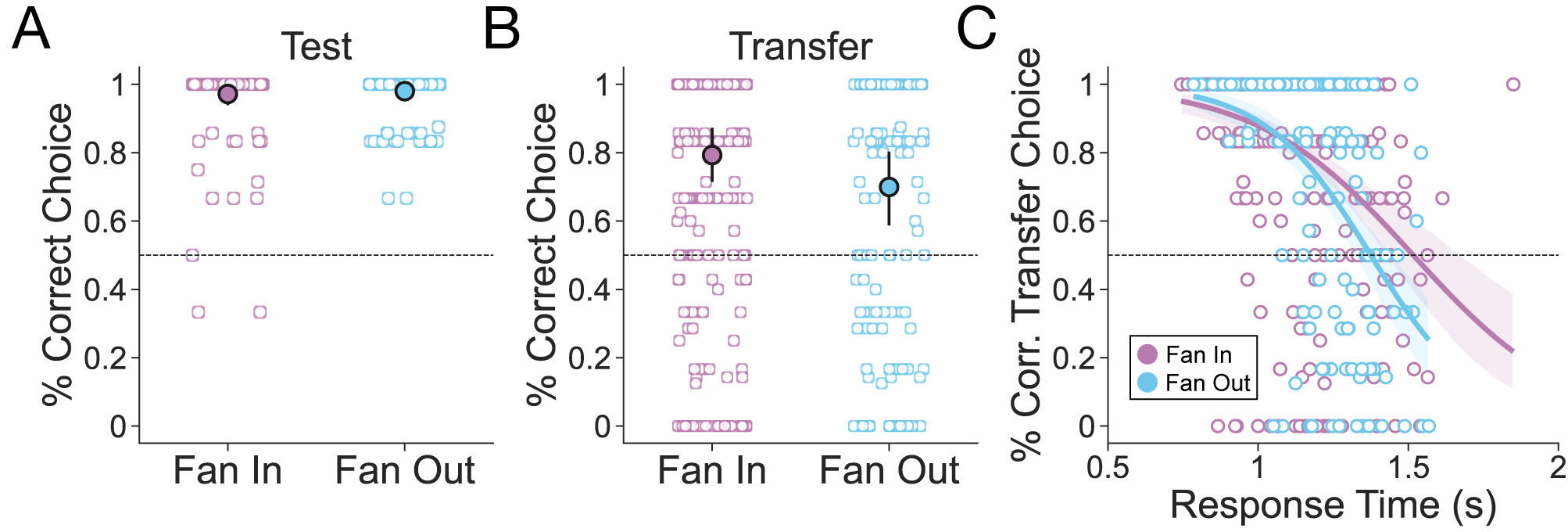
Participants successfully learned and transferred across both conditions, but the relationship between speed and accuracy differed across conditions. **A)** Test decisions (i.e. those between images that were directly associated with reward or neutral outcomes during reward learning) were highly accurate, reflecting successful learning for both conditions. **B)** Transfer decisions (i.e. those between images that were indirectly associated with reward or neutral outcomes via the stimulus learning phase) were also highly accurate, indicating successful inference for both conditions. Filled points represent group-level means whereas white points represent means for each pair of images seen by n=39 participants. Error bars are 95% confidence intervals. **C)** The relationship between the proportion of accurate transfer choices and reaction time for each image pair revealed that faster decisions were more accurate and that this relationship was stronger for the Fan Out condition, in which the structure was more amenable to proactive integration. Lines represent regression fits and bands represent 95% confidence intervals. Individual points represent means for each image pair. All visualizations show data at the stimuli level, and statistical analyses were conducted using mixed effects models that additionally assessed these effects within each participant while accounting for variation across participants.

Next, we examined participants’ transfer choices during the decision phase (**Figure 2B**). We found that participants tended to choose the antecedent image that was paired with the rewarded consequent image (β_’_ = 2.075, 95% *CI* = [1.283, 2.896]), indicating that most participants used memory to transfer value. There was no difference in transfer performance between Fan In and Fan Out choices (β*_conditions_* = 0.572, 95% *CI* = [−0.157, 1.284]), demonstrating that the manipulation of associative structure between conditions had no effect on the degree to which value was transferred.

Having established that participants infer the value of associated antecedent images in both conditions, we next sought to gain initial insights into *when* memories are accessed to support this value transfer. We aimed to differentiate between two possible strategies for inference, each occurring at different timepoints in our task: either proactively at reward learning or reactively at decision time. One hypothetical hallmark of proactive inference is that it should promote accuracy without the need for further memory retrieval of consequents at choice time, resulting in faster transfer decisions. Thus, if its deployment varies across stimuli, it predicts an unusual inverted speed-accuracy relationship whereby faster decisions tend also to be more accurate. In contrast, successful reactive inference by definition requires retrieving associations between memories at choice time, resulting in slower transfer decisions and (to the extent its deployment governs successful performance) a more typical relationship between slower decisions and higher accuracy.

Overall, we found that choices reflecting memory-based transfer were faster (β*_rt_* = −0.611, 95% *CI* = [−0.945, −0.287]; **Figure 2C**), suggesting that participants may have inferred proactively. In addition, this relationship was stronger in the Fan Out than the Fan In condition (β*_conditions_*:*_rt_* = −0.465, 95% *CI* = [−0.937, −0.017]), consistent with our expectation that the Fan In condition is less amenable to proactive inference. Together, these behavioral findings suggest that while proactive inference may be common in performance overall, reactive inference may have been more commonly observed in the Fan In than the Fan Out condition.

### Neural evidence for proactive and reactive inference and their modulation by retrieval competition

While examining participants’ choices allowed us to assess the different behavioral signatures of proactive and reactive inference, choice behavior alone cannot capture when exactly memories were accessed throughout the task. To gain further insight into when memories were recalled to support inference, we used fMRI to obtain a neural signature of memory reactivation at different timepoints in our task (**Figure 3A**). As in past work^11,12^, here we primarily interpret memory reactivation as a marker of inference, but note that another plausible role for memory reactivation may be to strengthen associations between individual memories^1,2^. To measure memory reactivation, we first used runs of fMRI data collected from the stimulus learning phase to train a classifier to distinguish between each image category: faces, scenes or objects. We then tested this classifier on activity from the reward learning and decision making phases, and assessed its ability to identify the category of the image that was presented to participants. As expected, voxels that differentiated accurately between categories were located primarily across the bilateral occipito-temporal cortex (**Figure 3B**). When tested on the reward learning and decision making phases, the classifier accurately differentiated each category from the others (Faces: β_0_ = 0.161, 95% *CI* = [0.134, 0.189]; Scenes: β_0_ = 0.151, 95% *CI* = [0.123, 0.180]; Objects: β_0_ = 0.066, 95% *CI* = [0.041, 0.093]; **Figure 3C**).

**Figure 3.**
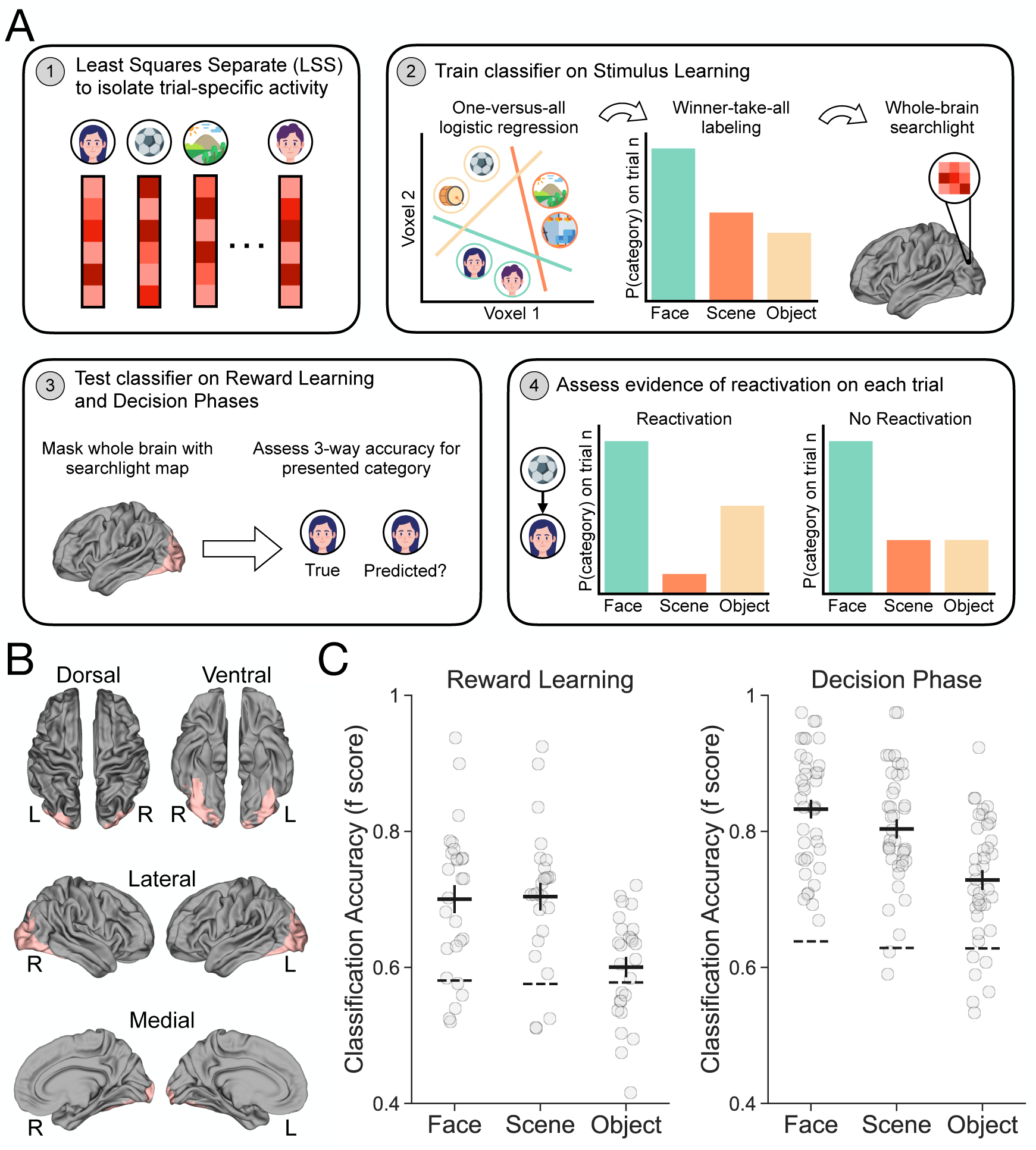
Multivariate pattern analysis methodology and decoding accuracy. **A)** MVPA analyses consisted of four primary steps. Step 1: Least Squares Separate^35^ was used to isolate a beta map for each trial and participant across all phases of the experiment. These betas were then used as input for the MVPA pipeline. Step 2: A searchlight analysis consisting of a one versus all three-way logistic regression was then used to identify voxels that could discriminate between all three categories during the stimulus learning phase. Step 3: Voxels identified during the previous step were then used to mask the whole brain during testing of the classifier on the reward learning and decision phases. Step 4: Evidence of reactivation on each trial was then assessed by ranking the individual category probabilities accordingly. **B)** Group-level whole-brain maps (FDR corrected; q<0.05) of voxels that discriminate between all three categories above chance. **C)** Classification accuracy for the decoding model trained on the stimulus learning phase and tested on the reward learning and decision phases. Accuracy is shown here as the weighted F-score. Points represent accuracy for each participant (n=39) and the thick line represents group-level average accuracy. Dotted lines represent the 95^th^ percentile of a permutation distribution over test category labels.

With a classifier in hand that could distinguish between each category based on BOLD activity patterns seen during the reward learning and decision phases, we were poised to assess the degree to which memories were reactivated for inference, and when. Specifically, to measure memory reactivation, we examined the individual category probabilities from the classifier on every trial, and identified those in which the probability of the *associated* image category (as opposed to the presented category) was particularly high (see **Methods**). This analysis allowed us to label every trial as one in which reactivation of the relevant associated category in memory was either likely or unlikely.

To determine whether memories were accessed in accordance with the patterns of inference we observed behaviorally, we focused on three main goals for the analyses. First, because participants’ choice behavior at transfer suggested a tradeoff between speed and accuracy most consistent with proactive inference, we sought to examine whether greater memory reactivation during the reward learning phase indeed results in more efficient (faster and more accurate) choices. Second, because we found that this effect was weaker during Fan In compared to Fan Out transfer choices (when there was relatively more retrieval competition between memories during reward learning and less during decision making), we sought to determine whether this behavioral shift was supported by different memory access patterns across conditions. Third, we predicted that it would be most strategic for participants to proactively infer prior to choice time for Fan Out trials, but to reactively infer at choice time for Fan In trials and therefore tested this by characterizing individual differences in memory access between participants.

To first examine whether memory access during reward learning leads to more efficient choices, we quantified the difference in memory reactivation during image viewing at reward learning and decision time. This yielded an index of proactive inference for each pair of images. We focused on the Fan Out condition because the design allowed us to measure reactivation for this condition at both of these time points (for the Fan In condition, the design only allows measuring reactivation at decision time; see **Methods**). When there was more evidence of proactive inference – i.e. when memory reactivation was greater at the time of reward learning relative to that of decision making – transfer choices were both more accurate (β_Δ_*_reactivation_* = 0.302, 95% *CI* = [0.0384, 0.593]) and marginally faster (β_Δ_*_reactivation_* = −37.902, 90% *CI* = [−75.273, −2.508], 95% *CI* = [−82.823, 3.180]; **Figure 4**). This result suggests that using memory to transfer value via proactive inference offers the advantage of more efficient choices in the future.

**Figure 4.**
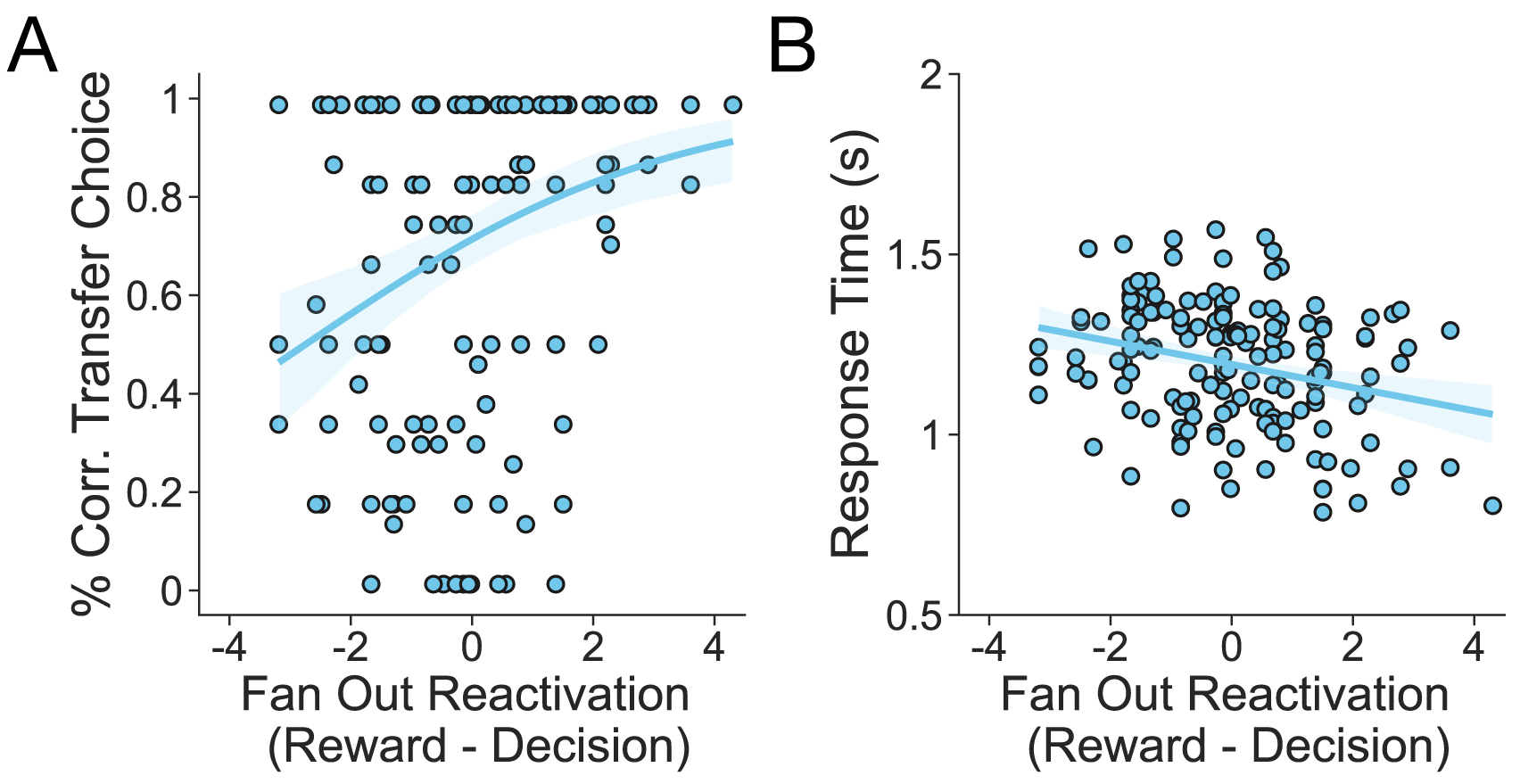
Proactive inference improves decision making ability. Greater memory reactivation at reward time relative to decision time – a marker of proactive inference – is associated with more effective transfer decisions. **A)** Correct transfer decisions were more likely for pairs with greater memory reactivation during reward learning relative to decision making. **B)** Response times were marginally faster for pairs with greater memory reactivation during reward learning relative to decision making. Points represent average performance for each image pair seen by participants. Lines represent regression fits and bands represent 95% confidence intervals. Visualizations show data at the stimuli level, and statistical analyses were conducted using mixed effects models that additionally assessed these effects within each participant while accounting for variation across participants.

We next examined whether the Fan In and Fan Out conditions affected memory access patterns, focusing on the time of choice because this was the timepoint at which we were best able to assess reactivation in both conditions (see **Methods**). In line with participants’ behavior, we found that during the decision phase, memories of rewarded consequent images were reactivated more frequently for Fan In than Fan Out transfer decisions (β*_condition_* = 0.119, 95% *CI* = [0.051, 0.184]; **Figure 5A**). This result indicates that our manipulation induced increased retrieval competition during Fan Out relative to Fan In transfer decisions. It further provides initial evidence that reactive inference may be more likely to occur when proactive inference is disadvantaged, although reduced Fan Out reactivation could also be consistent with accounts of reactive inference in which memories are retrieved in parallel (a point to which we return in the **Discussion**).

**Figure 5.**
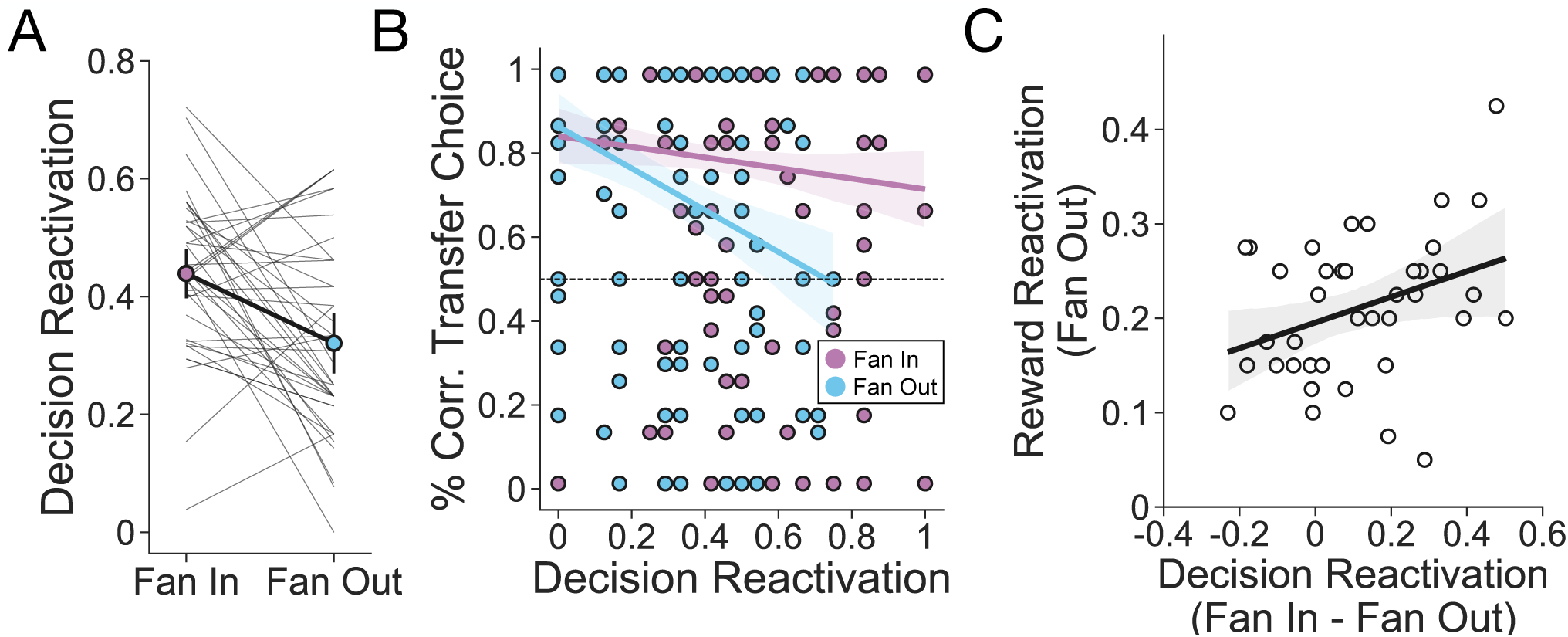
Reactive inference is more likely in the Fan In than Fan Out condition. **A)** Reactivation during the decision phase was greater for Fan In than Fan Out trials. Filled points represent group-level means, error bars are 95% confidence intervals, and thin lines represent individual participant slopes (n=39). **B)** Greater memory reactivation at decision time, a marker of reactive inference, is associated with less effective transfer decisions for Fan Out but not Fan In image pairs. Points represent average performance for each image pair seen by participants. Lines represent regression fits and bands represent 95% confidence intervals. **C)** Participants who showed greater reactivation for Fan In relative to Fan Out trials during decision making also preferentially reactivated more for Fan Out trials during reward learning. Points represent individual participant means, the line represents a linear regression fit, and the band represents a 95% confidence interval.

To further investigate the possibility that reactive inference is more likely when proactive inference is relatively less advantageous, we examined the relationship between decision-time memory reactivation and behavioral performance. The behavioral findings showed that transfer choices were both slower and less successful in the Fan Out relative to the Fan In condition (**Figure 2C**). This effect may reflect the fact that, due to competition, proactive inference is easier and reactive inference is correspondingly harder, making it less likely to be successful in the Fan Out condition. We therefore predicted that the neural measure of memory reactivation at decision time should likewise be associated with less successful value transfer in the Fan Out condition. Indeed, we found that Fan Out transfer decisions were less accurate when antecedent memories were reactivated at decision time (β*_reactivation_* = −0.300, 95% *CI* = [−0.625, −0.001]; **Figure 5B**). Further, no such effect was found in the Fan In condition (β*_reactivation_* = −0.086, 95% *CI* = [−0.255, 0.075]; **Supplementary** Figure 1). This result lends additional support to the interpretation that the manipulation of associative structure increased participants’ relative use of reactive inference in the Fan In condition.

Finally, we assessed the idea that it would be strategic to proactively infer prior to choice time for Fan Out trials, and to reactively infer at choice time for Fan In trials. We examined whether individuals who tend to reactivate memories more for Fan In relative to Fan Out trials at decision time also reactivated memories more for Fan Out trials during the reward learning phase. That is, we asked whether participants’ ability to appropriately deploy one of these strategies also predicted appropriate deployment of the other. We found that this was indeed the case— participants who reactivated memories more for Fan In transfer decisions relative to Fan Out transfer decisions also reactivated memories for Fan Out stimuli at reward learning (β_Δ_*_reactivation_* = 0.027, 95% *CI* = [0.003, 0.050]; **Figure 5C**). This result suggests that those participants who were most sensitive to the presence of retrieval competition at either timepoint strategically modulated when they accessed their memories to perform inference.

## Discussion

Research on sequential decision making has found that the process of linking memories to support inference is well described by theories of reinforcement learning that leverage an internal model to guide choice^4–6,9,10,18,19^. Numerous studies have shown that memory-based inference can occur at a number of different timepoints relative to the moment a decision is made^10–12,21,22,25,26,36,37^. However, the conditions that lead some memories to be accessed later than others have remained unclear. Here we developed a task to directly test multiple hypotheses about the purpose and adaptability of memory access in inference. Using fMRI to decode patterns of BOLD response unique to the categories of images in memory, we found that participants primarily accessed memories proactively, but this pattern was also sensitive to the situation: when a choice option had multiple past associations, participants were more likely to defer inferring relationships between stimuli and outcomes until decisions were made. This finding suggests that the presence of competition between associations in memory makes their retrieval for use in inference less likely, and runs counter to alternate possibilities in which the opposite may have been true (e.g., if memory reactivation is primarily driven by the imperative to associate reward with related stimuli, one may expect relatively more reactivation for the Fan In condition during reward learning and for the Fan Out condition during choice). We also found neural and behavioral evidence that reinstating memories prior to decision making facilitates faster and more accurate inference, suggesting that it is adaptive to plan in advance when possible. Together, these results indicate that the brain judiciously conducts proactive inference, accessing memories proactively in conditions when this is most favorable.

These findings add empirical support to predictions from a recent rational account of when each of these forms of inference is most useful for decision making^15^. Specifically, Mattar and Daw (2018) theorized that memories that are particularly likely to increase future expected reward will be prioritized for reinstatement during inference and planning. Formally, they proposed that the expected utility of accessing a past experience can be decomposed into the product of two terms: need and gain. Need quantifies how likely an experience is to be encountered again, and gain captures how much reward is expected from improved decisions if that experience is reinstated. A critical feature of this model is that when the need term dominates, memories tend to be accessed reactively at choice time, but if instead the gain term dominates, memories tend to be accessed proactively following the receipt of reward. The present findings generally support this theory. In particular, gain increases for an antecedent when choices fan out, favoring proactive memory access, while need increases for consequents, promoting reactive choice-time memory access, as they fan in. Thus, antecedents that are associated with many consequents (i.e. that fan out) are more likely to be reinstated upon learning that a consequent is rewarded, because there is much to gain from updating future decisions made upon future encounters with the antecedent. Likewise, antecedents which deterministically lead to a single consequent (i.e. that fan in) imply greater need for that consequent, and are more likely to be reinstated at decision time. Importantly, while our findings are consistent with this theory, they were also designed to be predicted by more intuitive, qualitative reasoning about the degree of competition among different memories, and so go beyond any single theory of prioritization for memory access.

In addition to findings from sensory preconditioning demonstrating that humans use memories for inference^11,12^, a number of other studies have shown that memory-based inference may also take place offline, during periods of rest or sleep before choice. This approach is advantageous because it offloads computation to otherwise unoccupied time. In humans, fMRI research has revealed that memories are reactivated during periods of rest following reward^23,24^ and that this reinstatement can enhance subsequent memory performance^38,39^. Importantly, such offline replay of past memories during rest has been shown to facilitate later integrative decisions^25,26^. Parallel work in rodents has demonstrated that hippocampal replay of previously experienced spatial trajectories is observed during rest and sleep^40–42^, and that rewarded locations are replayed more frequently^28^. These results indicate another way in which inferences may be drawn offline, well before constituent memories are needed for choice. An important direction for future work will be to see if rational considerations, such as sensitivity to competition between memories, also affect the likelihood, or targets of, offline inference.

A separate important open question regards the details of how associations between memories are represented in the brain. In other words, what is the nature of the internal model? Computational work on reinforcement learning has identified multiple candidate algorithms that may give rise to the effects reported here. Broadly, these theories posit that agents come to represent associations between states in an internal model, and then, using this model, simulate experiences to discover the consequences of new actions. The process of simulating potential actions can occur in either a forward or backwards manner, and can be based upon internal models with different representational forms. For example, in RL algorithms that employ a full world model, forward simulation is accomplished by adding up expected immediate rewards over some explicit future trajectory (rolled out over a series of one-step associations), while backwards simulation can occur by propagating value information from a destination state to a series of predecessors^43,44^. Other algorithms, such as successor^17^ and predecessor representations, learn temporally abstract state relationships that are aggregated over multiple timesteps, and can be similarly used to compute which states typically follow or precede the present, respectively. Our findings are consistent with either of these frameworks.

Moreover, our experimental design is not well positioned to differentiate between them. Using tasks in which future outcomes are separated from the present by multiple steps in space or time, much work has found that people may encode both one-step transitions between states and aggregate summaries of these relationships over multiple timesteps in the form of a successor representation^18,19,37,49^. While in such multi-step tasks these approaches often lead to substantially different internal models (and specifically, more efficient noniterative inference for successor representations), on tasks that involve only a one-step relationship between an antecedent and a consequent (such as the task we used here), their internal models are roughly equivalent. For this reason, the results of the current study have no bearing on this distinction. With that said, it is possible that these approaches may differ in their computational costs even in one-step tasks: implementations of the successor representation often assume that all successors are visited in parallel (as by a dot product) whereas those using a full transition model often employ serial rollouts or tree search. Regardless of the form of model participants relied upon to complete the task and the particular steps involved in using it for evaluation, our results are consistent with the idea that proactive inference yields benefits by eliminating the need to retrieve associations between memories at choice time.

Relatedly, in our study we are unable to isolate *how* people may retrieve memories from their internal models. While it is the case that algorithms employing a successor representation typically retrieve states in parallel and those incorporating a full transition model typically do so serially, several formulations exist in which the opposite is true^50–53^. Both of these forms of retrieval may have been used to support transfer choices in our task, and we are unable to clearly differentiate between them in the present work. Although our results are broadly consistent with serial retrieval, reduced reactivation of the rewarded consequent image relative to the other associated consequent image during Fan Out transfer choices (**Figure 5A**) is also consistent with inference algorithms that retrieve in parallel. This is because parallel retrieval would predict equal reactivation of both consequent stimuli. We note, however, that our other findings are unlikely to be explained by such an account. Determining both the form of representation people use for proactive and reactive inference and how memories are accessed to support inference more broadly remain questions for future research. This is particularly important because the advantages offered by computing value proactively may be offset by using a successor representation for reactive inference in environments with a particularly large temporal horizon or where the reward values of states may change.

In connection with these points, recent behavioral work in humans has also shown that efficient one-step predictive representations are used for both forwards prediction at decision time and also backwards prediction in a manner similar to the proactive inference strategy we measured here^54^. In particular, this study demonstrated that such a strategy is relied upon more often in environments where the number of states that follow a starting state outnumber those that precede a rewarded state. Using a similar manipulation coupled with direct assays of strategy use, our results provide convergent evidence for this idea. Our study further enhances understanding of proactive and reactive approaches to inference by grounding each of these strategies in the mechanisms of memory.

A separate avenue for future study that we did not touch upon here involves the role of dopamine in supporting the integration of memories with reward to guide behavior. Although the dopaminergic system has traditionally been thought to support habitual learning from direct experience, recent results suggest that dopamine may also support integrative evaluations of actions through the flexible combination of past experience^55,56^. Our task may provide an opportunity to further elucidate the role of dopamine in this process. Despite being solved in different ways, both of the conditions in our task are dependent upon the flexible expression of knowledge about stimulus associations. Therefore, if dopamine is necessary for the acquisition of model-based associations, as has been recently suggested^56^, we expect it to be involved in both conditions equally. This prediction could, for example, be tested by examining how integrative choice behavior in the present task is affected by dopamine depletion in Parkinson’s disease.

Other open questions remain about the precise role of memory reactivation in designs such as ours. Following prior research^11,12^, we used stimuli from specific visual categories to measure category-specific BOLD activity as a proxy for memory reactivation. Here, as in this past work, we interpreted memory reactivation in our design as a sign of memory-based inference, or retrieval to transfer reward information across associated states. But another role of reactivation may be to strengthen previously learned associations between individual memories (e.g., to build or update a successor representation as in the Dyna-SR algorithm rather than transferring reward associations as heretofore assumed^36,37^). It is possible that this mechanism may contribute to the effects reported here; for example, reactivating memories prior to choice (during our reward phase) may prevent forgetting (e.g., by strengthening or updating the associative model), leading to improved inferences in the future (e.g., by manifesting here as changes in reaction times or neural measures of retrieval during the transfer phase). Our study cannot fully rule out this possibility, particularly in how it may contribute to the improved benefits of proactive inference at decision time that we measured for Fan Out stimuli (**Figure 4**). However, past work on the sensory preconditioning task suggests that reactivation during the reward learning phase likely measures proactive inference, at least in part. Specifically, Kurth-Nelson et al., 2015 found that successful transfer decisions were associated with greater memory reactivation at outcome time (i.e., during the presentation of reward information). This finding appears to be best explained by proactive inference about reward, which predicts that credit should be assigned to the antecedent at this moment. Outside of this evidence, it is also important to note that the alternative still corresponds with our general framework: associations between memories can be strengthened ahead of time, providing future benefits, or during choices themselves, leading to similar tradeoffs in speed and accuracy. In fact, there are several theoretical accounts in which replay has this effect^36,37^. Disentangling these possibilities remains a critical goal for future work. One approach for future studies may be, for example, to include more explicit measures of memory for each stimulus association.

Separately, one shortcoming of our study was that, due to our design, we were unable to isolate memory reactivation when consequent images from the Fan In condition were presented during reward learning. In practice, this limited our contrasts between conditions to decision time and our contrasts between timepoints to the Fan Out condition. This was because our metric of memory reactivation was conservative in the sense of being selective to the specific relevant candidate for classification. In particular, in addition to the category actually present on the screen being most strongly decoded, we required that the relevant associate be more strongly activated than the irrelevant foil to declare reactivation successful. However, at reward time in the Fan In condition, both categories are relevant associates, so this comparison was not possible. One possibility to skirt this issue in future work may be to present images of a fourth entirely unrelated category. We did not pursue this direction in the present study to minimize the complexity of the design. Future complementary work may explore these issues in more depth in order to allow for cleaner measurement of reactivation when antecedent images fan in during reward learning.

In conclusion, we have demonstrated that the statistical structure of training experience impacts whether inference from memory occurs before or during decision making. This finding suggests that standard prospective inference is not unique, but is instead one of a general set of computations that access memory at different times. Together, these findings further help to explain why different studies have observed memory integration to support choice at different times, and suggest that different inference strategies may be recruited depending on their efficacy for the task at hand.

## Methods

### Participants

A total of 40 participants (19 M, 21 F) between the ages of 18 – 35 were recruited from the Columbia University community. Participants were right-handed, had normal or corrected-to-normal vision, took no psychiatric medication, and had no diagnosis of psychological disorders. One participant was removed from the analyses due to both failing to understand the instructions of the task and missing responses on over half of the decision trials. The remaining 39 participants had a mean age of 21.9 with a range of 19-35 and were included in the reported sample. No statistical method was used to predetermine sample size. Informed consent was obtained at the beginning of the session and all experimental procedures were approved by the Columbia University Institutional Review Board.

### Experimental Task

Participants completed a three-part associative learning task while undergoing an fMRI scan. In the first phase of the experiment, *stimulus learning*, participants were tasked with learning pairs of images presented one at a time. Each trial consisted of a single image (A; 1.5s), followed by a interstimulus interval in which a fixation cross was displayed (exponentially jittered with mean=3s, min=0.5s, max=12s), followed by another image (B; 1.5s), and finally an intertrial interval in which another fixation cross was displayed (exponentially jittered with mean=3s, min=0.5s, max=12s). In order to ensure that participants were paying attention, they were asked to press a button box with their index finger for the first image and with the middle finger for the second image in a pair. Participants were shown 16 different pairs of images 5 times each for a total of 80 trials. Trials were spread across two runs of 40 trials each. Images came from one of three categories, either a face, a scene, or an object. In the second phase of the experiment, *reward learning,* participants were tasked with learning that a subset of B images from the stimulus learning phase led deterministically to reward, while another subset of images led deterministically to a neutral outcome. Each trial consisted of a single image (1.5s), followed by an interstimulus interval in which a fixation cross was displayed (2s), followed by the outcome (either a dollar bill or a gray rectangle; 1.5s), and then finally an intertrial interval (exponentially jittered with mean=2.5s, min=0.5s, max=10s). Participants were told to withhold a response for the image and to respond with their index finger when a dollar was shown and with their middle finger when a gray rectangle was shown. Participants saw each of 8 images 10 times for a total of 80 trials. Trials were spread across two runs of 40 trials each. During the third and final phase of the experiment, the *decision phase*, participants were tasked with deciding between two images of the same category (either A v. A or B v. B) presented on the screen simultaneously. Each trial consisted of a choice (max=2s), a confirmation in which a green rectangle appeared around their choice (2s-reaction time), and then an intertrial interval (exponentially jittered with mean=2.5s, min=0.5s, max=10s). Participants pressed with their index finger to choose the image on the left hand side of the screen and with their middle finder to choose the image on the right hand side of the screen. Participants made 78 choices across a single run of this phase. Interstimulus intervals and trial ordering was optimized to minimize the correlation between events throughout each phase of the task.

The pairs of stimuli presented throughout the experiment fell into one of two conditions that were unknown to participants: Fan Out and Fan In trials. Fan Out trials consisted of one A image that could be followed by either of two B images, while Fan In trials consisted of either of two A images followed by one B image. During stimulus learning, eight pairs of images fanned in, while another eight fanned out. Of the eight pairs from each condition, there were two pairs of images for each of four possible combinations (e.g. Fan In: A1-B1; A2-B1; A4-B4; A5-B4; Fan Out: A3-B2; A3-B3; A6-B5; A6-B6). During reward learning, four B images from each condition were shown (e.g. Fan In: B2 x2; B5 x2; Fan Out: B1 x2; B4 x2) such that two from each condition were paired with reward (e.g. Fan In: B1; Fan Out: B2) and two were paired with a neutral outcome (e.g. Fan In: B4; Fan Out: B5). B3 and B6 stimuli were not shown during this phase and were not associated with any outcome. Finally, during the decision phase, participants made choices between B images that had been directly associated with a reward or neutral outcome (*test* choices) and between A images that had been indirectly associated with these outcomes (*transfer* choices). Test (e.g. Fan In: B1 v B4; Fan Out: B2 v B5) and transfer (e.g. Fan In: A1 v A4; A2 v A5; Fan Out: A3 v A6) choices were made between images from the same condition, and never between images from different conditions.

Participants were told prior to starting the task that they would need to use the associations they learned throughout the first two phases of the experiment in order to make choices in the final phase. They were given a cover story to aid their learning throughout the task. Specifically, participants were told that they were a photographer visiting a new city and would be taking different buses to different locations. At each location, they would be shown a picture they had taken there, and the purpose of the first phase was to learn which photos were taken along each bus route. Then, during the reward learning phase, participants were told that they had returned from their trip and had sent their photos to clients for potential purchase. They were then shown which photos had been purchased and which had not, and their goal was to learn this information. Lastly, during the decision phase, participants were told that they were planning a new trip to the city and were tasked with deciding between bus routes (represented by photos taken on each route) that would take them to locations where they had taken photos their clients purchased. Participants were instructed to use what they had learned (i.e. which photos were taken along the same route and which were or were not purchased) to inform their choices.

### MRI Acquisition

MRI data were collected on a 3 T Siemens Magnetom Prisma scanner with a 64-channel head coil. Functional images were acquired using a multiband echo-planer imaging (EPI) sequence (repetition time = 1.5s, echo time = 30ms, flip angle = 65°, acceleration factor = 3, voxel size = 2 mm iso, acquisition matrix 96 x 96). Sixty nine oblique axial slices (14° transverse to coronal) were acquired in an interleaved order and spaced 2mm to achieve full brain coverage. Whole-brain high resolution (1 mm iso) T1-weighted structural images were acquired with a magnetization-prepared rapid acquisition gradient-echo (MPRAGE) sequence. Field maps consisting of 69 oblique axial slices (2 mm isotropic) were collected to aid registration.

### Imaging Data Preprocessing

Results included in this manuscript come from preprocessing performed using *fMRIPrep* 20.2.6, which is based on *Nipype* 1.7.0.^57^

### Anatomical Data Preprocessing

Each participant’s T1-weighted (T1w) image was corrected for intensity non-uniformity (INU) with N4BiasFieldCorrection^58^, distributed with ANTs 2.3.3^59^ and used as a reference image throughout the workflow. The reference image was then skull-stripped with a *Nipype* implementation of the antsBrainExtraction.sh workflow (from ANTs), using OASIS30ANTs as target template. Brain tissue segmentation of cerebrospinal fluid (CSF), white-matter (WM) and gray-matter (GM) was performed on the brain-extracted T1w using fast^60^ (FSL 5.0.9). Volume-based spatial normalization to the *ICBM 152 Nonlinear Asymmetrical template version 2009c* (MNI152NLin2009cAsym) standard space was performed through nonlinear registration with antsRegistration (ANTs 2.3.3), using brain-extracted versions of both the T1w reference and the T1w template images.

### Functional Data Preprocessing

For each of the 5 BOLD runs per participant (two stimulus learning runs, two reward learning runs, and one choice run), the following preprocessing was performed. First, a reference volume and its skull-stripped version were generated using a custom methodology of *fMRIPrep*. A B0-nonuniformity map (or *fieldmap*) was estimated based on two (or more) echo-planar imaging (EPI) references with opposing phase-encoding directions, with 3dQwarp^61^ (AFNI 20160207). Based on the estimated susceptibility distortion, a corrected EPI reference was calculated for a more accurate co-registration with the anatomical reference. The BOLD reference was then co-registered to the T1w reference using bbregister (FreeSurfer) which implements boundary-based registration^62^. Co-registration was configured with six degrees of freedom. Head-motion parameters with respect to the BOLD reference (transformation matrices, and six corresponding rotation and translation parameters) were estimated before any spatiotemporal filtering using mcflirt^63^ (FSL 5.0.9). BOLD runs were slice-time corrected to 0.708s (0.5 of slice acquisition range 0s-1.42s) using 3dTshift from AFNI 20160207^61^. The BOLD time-series (including slice-timing correction when applied) were resampled onto their original, native space by applying a single, composite transform to correct for head-motion and susceptibility distortions. The BOLD time-series were resampled into standard space, generating a preprocessed BOLD run in MNI152NLin2009cAsym space. First, a reference volume and its skull-stripped version were generated using a custom methodology of *fMRIPrep*. Several confounding time-series were calculated based on the preprocessed BOLD: framewise displacement (FD), DVARS and three region-wise global signals. FD was computed using two formulations following Power (absolute sum of relative motions)^64^ and Jenkinson (relative root mean square displacement between affines)^63^. FD and DVARS are calculated for each functional run, both using their implementations in *Nipype*. The three global signals are extracted within the CSF, the WM, and the whole-brain masks. The head-motion estimates calculated in the correction step were also placed within the corresponding confounds file. The confound time series derived from head motion estimates and global signals were expanded with the inclusion of temporal derivatives and quadratic terms for each^65^. Frames that exceeded a threshold of 0.5 mm FD or 1.5 standardized DVARS were annotated as motion outliers. All resamplings can be performed with a single interpolation step by composing all the pertinent transformations (i.e. head-motion transform matrices, susceptibility distortion correction when available, and co-registrations to anatomical and output spaces). Gridded (volumetric) resamplings were performed using antsApplyTransforms (ANTs), configured with Lanczos interpolation to minimize the smoothing effects of other kernels^66^. Preprocessed data were lastly smoothed using a Gaussian kernel with a FWHM of 6.0mm, masked, and mean-scaled over time.

### Functional Imaging Data Analysis

#### Beta Series Modeling

Least squares separate (LSS) models were generated for each event (presentation of a category image) in each task following the method described in Turner et al., 2012^35^ using *Nistats 0.0.1b2*. For each trial, preprocessed data were subjected to a general linear model in which the trial was modeled in its own regressor, while all other trials from that condition were modeled in a second regressor, and other conditions were modeled in their own regressors. Each condition regressor was convolved with the *glover* hemodynamic response function for the model. In addition to condition regressors, 36 nuisance regressors were included in each model consisting of two physiological time series (the mean WM and CSF signals), the global signal, six head-motion parameters, their derivatives, quadratic terms, and squares of derivatives. Spike regression was additionally performed by including a regressor for each motion outlier identified in each run, as in Satterthwaite et al., 2013^65^. A high-pass filter of 0.0078125 Hz, implemented using a cosine drift model, was also included in each model and AR(1) prewhitening was applied to each model to account for temporal autocorrelation. After fitting each model, the parameter estimate (i.e., beta) map associated with the target trial’s regressor was retained and used for further analysis. Modeling was performed using *NiBetaSeries* 0.6.0^67^ which is based on *Nipype* 1.4.2.^57^ Beta maps for image presentation events, separated by category, for the stimulus learning and reward learning phases and for decisions between images, again separated by category, were used in subsequent analyses.

#### Multivariate Pattern Decoding Analysis

Beta maps from each trial were next used for multivariate pattern analysis. First, a searchlight classification analysis was conducted for each participant. In brief, a three-way one versus all logistic regression classifier was trained to distinguish categories using leave-one-run-out cross validation from runs of the stimulus learning task. We used winner-take-all labeling to determine the classified label from each trial: the category resulting in the highest probability from the one versus all classification procedure on a given trial was selected as the predicted label for that trial. Input data were selected using a spherical searchlight (radius = 2 voxels) moved around the whole brain. Although the experimental design leads the class labels for each category to be imbalanced during the stimulus learning phase (i.e. one label always has twice as many occurrences as the other two), we dealt with this label imbalance in two ways. First, the class weights applied to each category by the classifier were determined using the ‘balanced’ keyword in *sklearn*^68^ such that the weights were the number of samples divided by the number of labels (3) multiplied by the total number of occurrences of each label. Second, our metric of performance was the weighted-F1 score, which is the harmonic mean of precision and recall. Each of these methods are commonly used in the machine learning literature to deal with class imbalance in training data. For each searchlight sphere, we additionally computed chance performance via a permutation test: labels were shuffled 1000 times and the weighted F1-score resulting from each of these permutations was computed. Chance classification performance was then calculated as the 95^th^ percentile of the F1-score permutation distribution. For each voxel, we then subtracted chance level performance from the classification accuracy to produce a map of corrected classification performance for each participant. Finally, an FDR-corrected (q<0.05) group-level map over all individual participant difference maps was created.

Following classifier training on the stimulus learning phase, we then tested the classifier on runs from both the reward learning and decision phases. Functional data from each participant on each of these phases of the experiment was first masked using the group-level searchlight map produced from the previously described procedure. The three-way logistic regression classifier was then re-trained on both runs of the stimulus learning phase, using only these voxels, and then tested separately on the reward learning and decision phases. L1-regularization was used to reduce overfitting in this procedure. We again used the weighted F1-score as our accuracy metric, and the 95^th^ percentile of the permutation distribution as our measure of chance classifier performance.

Finally, to address our primary question, we created an index of memory reactivation from the classifier. Specifically, for each trial, we extracted the probability that the classifier assigned to each category label. A trial was then considered a trial on which memory reactivation occurred if the following criteria were met: i) the true category label was assigned the highest probability by the classifier and ii) the associated category was assigned the second highest probability by the classifier. If these criteria were met, the trial was assigned a one and, if not, a zero. Our logic for using this criteria was conservative: we reasoned that the classifier should always assign the highest probability to the category represented by the image that is presently shown on the screen. Because, by definition, both off-screen categories were candidates for association when presented as part of Fan In trials during the reward learning phase, we were unable to calculate a reactivation score for these trials. We were further limited in our ability to compare reactivation across phases because the classifier was more accurate at identifying category images presented during the decision phase than during the reward learning phase. This is problematic because lower classification accuracy causes lower reactivation scores because fewer trials satisfy the criteria outlined above. We were, however, able to investigate individual differences in reactivation for Fan Out trials between phases by accounting for this difference in classification performance by z-scoring reactivation scores within each phase, as this removes group-level differences while leaving individual differences intact. These standardized reactivation scores were used only for analyses involving comparison between phases of the experiment.

### Regression Analyses

Unless otherwise noted, parameters for all regression models described here were estimated using hierarchical Bayesian inference such that group-level priors were used to regularize participant-level estimates. The joint posterior was approximated using No-U-Turn Sampling^69^ as implemented in stan. Four chains with 2000 samples (1000 discarded as burn-in) were run for a total of 4000 posterior samples per model. Chain convergence was determined by ensuring that the Gelman-Rubin statistic *R*^7^ was close to 1. Default weakly-informative priors implemented in the *rstanarm*^70^ package were used for each regression model. For all models, fixed effects are reported in the text as the mean of each parameter’s marginal posterior distribution alongside 95% or 90% credible intervals, which indicate where that percentage of the posterior density falls. Parameter values outside of this range are unlikely given the model, data, and priors. Thus, if the range of likely values does not include zero, we conclude that a meaningful effect was observed.

We first assessed choice performance on the decision phase of the task. For each participant. *s* and trial *t*, a mixed effects logistic regression was used to predict if the correct image was chosen:

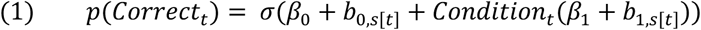

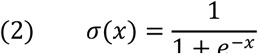

where *Correct* was equal to 1 if the participant chose either the image directly associated with reward (in the case of test trials) or the image indirectly associated with reward (in the case of transfer trials), and *Condition* was a categorical variable coded as 0.5 for Fan In trials and –0.5 for Fan Out trials. This model was fit separately for test and transfer choices.

We also assessed the relationship between response time and accuracy during transfer choices using the following mixed effects logistic regression, which included an additional main effect of response time as well the interaction between response time and condition:

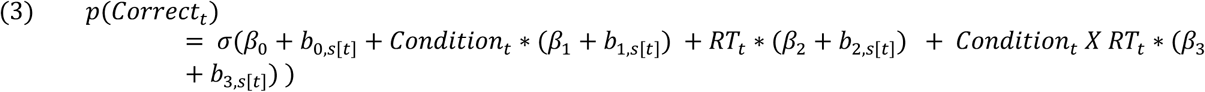

where *RT* was the response time on each transfer choice trial.

We determined the ability of the trained MVPA classifier to distinguish each category label from chance using the following mixed effects linear regression:

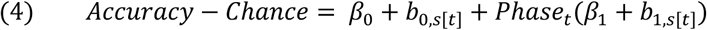

where *Accuracy* − *C*ℎ*ance* was the 95^th^ percentile of the permutation distribution subtracted from classification accuracy, and *P*ℎ*ase* was a categorical variable coded as 0.5 for the decision phase and –0.5 for the reward learning phase. This model was fit separately for each category (face, scene and object).

Another set of models was fit to assess the relationship between memory reactivation and transfer choice behavior. Analyses were conducted on the average reactivation level of each stimulus. In order to assess effects of reactivation on transfer accuracy for each stimulus, *i*, accuracy was first transformed^71^ to ensure that all responses fell within the interval (0,1):

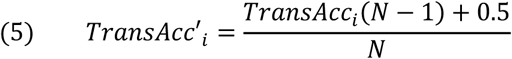

where *TransAcc* was participants’ average transfer accuracy for each consequent stimulus and *N* was the sample size (39). We first examined the effect of (z-scored) differences in reactivation between the reward learning and decision phases for each associated antecedent-consequent pair of Fan Out stimuli on transfer accuracy. To do so, we fit a mixed effects beta regression:

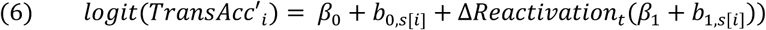

where Δ*Reactivation* is the difference in memory reactivation between reward learning and the decision phase for each pair. Similar beta regressions were used to assess effects of memory reactivation during the decision phase for Fan In and Fan Out consequent stimuli, separately. To assess effects on choice transfer response time, linear mixed effects regressions with the same predictors were used instead.

We additionally assessed how memory reactivation differed for each condition (Fan In or Fan Out) during the decision phase. We performed this analysis using the following mixed effects linear regression:

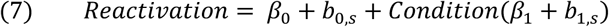

where *Reactivation* was memory reactivation during the decision phase for each participant and condition and *Condition* was coded identically to the models described above.

Lastly, we examined individual differences in strategy usage by comparing our reactivation measures across phases of the task. Specifically, we fit a simple linear regression predicting each participants’ average level of memory reactivation for Fan Out during reward learning from their difference in memory reactivation during the decision phase.

## Data Availability

The data that support the findings of this study are available in GIN with identifier: 10.12751/g-node.ee5wx3

## Code Availability

The code used to generate the results of this study are available as a CodeOcean capsule with identifier: 10.24433/CO.2559896.v1

## Acknowledgements

The authors thank members of the Shohamy Lab for insightful discussion and Paul Sharp and the other anonymous reviewer for their feedback and comments. Support was provided by the Kavli Foundation (D.S.), NSF GRFP (J.N.; award #1644869), the NSF (D.S., N.D.D.; award #1822619), the NIMH/NIH (D.S., N.D.D.; award #MH121093) and the Templeton Foundation (D.S. grant # 60844). Figures 1 and 3 use images created by Creartive, Freepik, and Paul J on Flaticon.com.

## Author Contributions

Conceptualization, J.N., N.D.D., D.S.; Methodology, J.N., N.D.D., D.S.; Software, J.N.; Formal Analysis, J.N.; Investigation, J.N.; Writing – Original Draft, J.N.; Writing – Review & Editing, N.D.D, D.S.; Visualization, J.N., N.D.D., D.S.; Supervision, D.S.; Funding Acquisition, N.D.D., D.S.

