## Supplemental Information for "Proactive and reactive construction of memory-based preferences"

### Supplementary Information

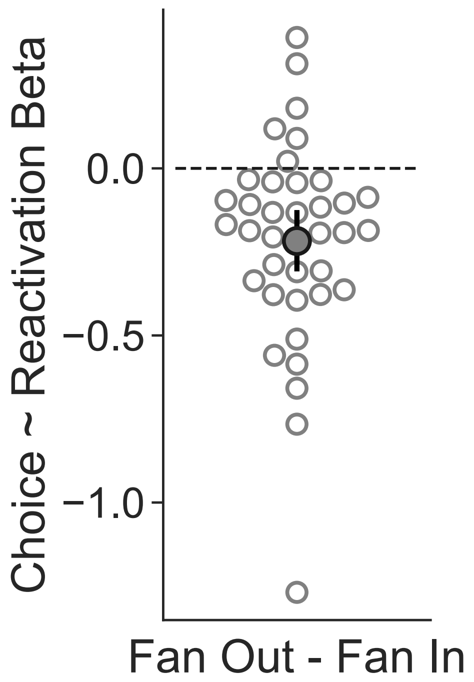


**Supplementary Figure 1.** As shown in Figure 4C, greater memory reactivation at decision time is associated with less effective transfer decisions for Fan In but not Fan Out image pairs (n=39 participants). Shown here is this effect, the difference in slopes, for every participant and at the group-level. Participants demonstrate this relationship more for Fan Out than Fan In decisions ($\beta_0 = -0.215$, $95\%\ CI = [-0.312, -0.118]$), as indicated by comparing their random slopes. The filled point represents the group-average difference in slopes, whereas empty points represent individual slope differences.

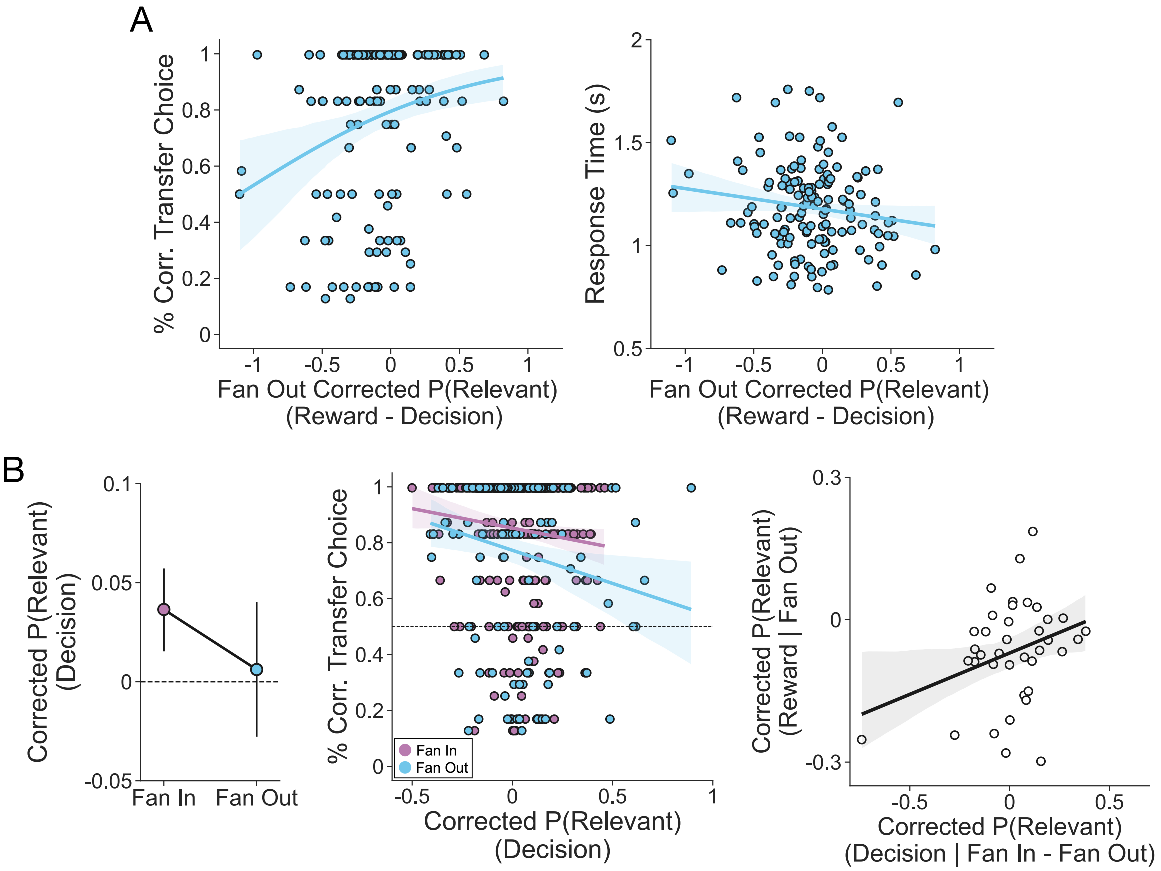


**Supplementary Figure 2.** All reactivation-based analyses repeated using a measure based directly on the classifier probabilities upon which the reactivation score is built. We repeated our analyses using the classifier probability for the associated category that is required for inference in each condition (labeled here as “relevant”). We subtracted from this value the probability of the category that was not needed for inference, which provides a baseline level of classifier probability related to noise on each trial. Using this measure (Corrected P(Relevant)), we found the same patterns for all results related to Figure 4 (A) and Figure 5 (B).

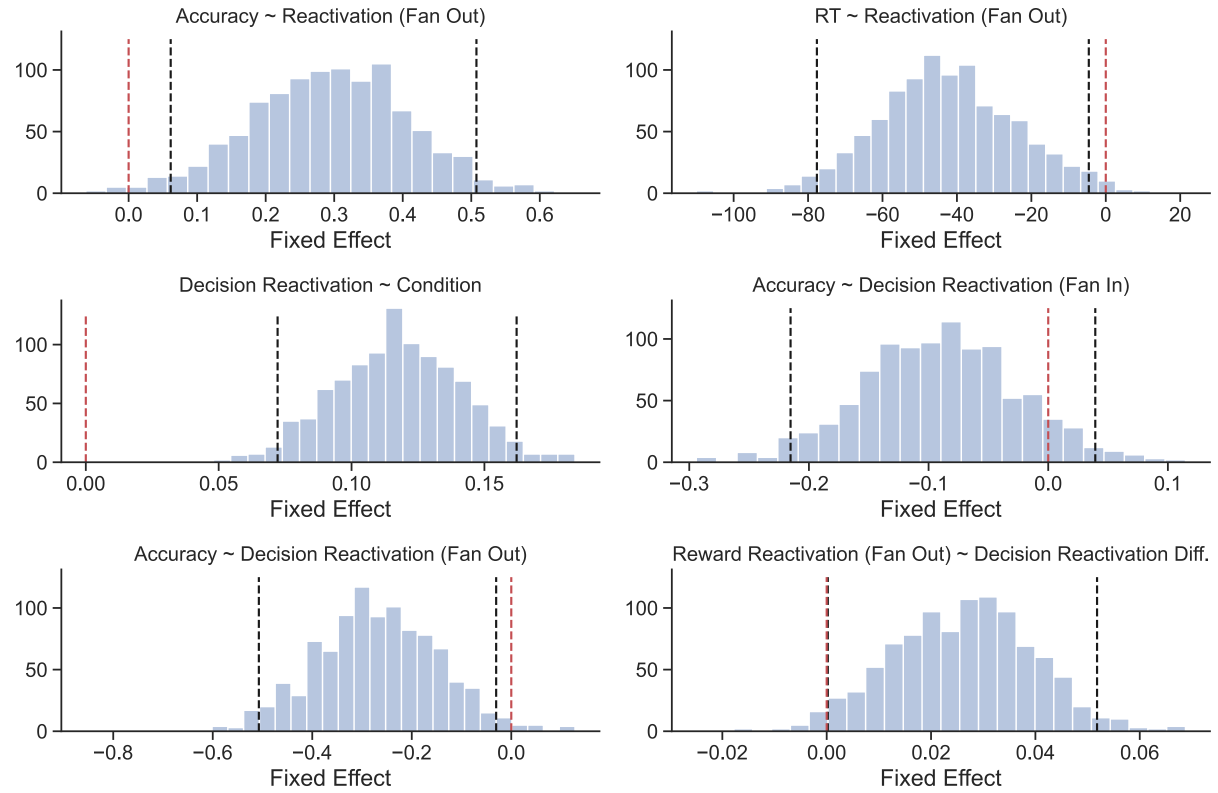


**Supplementary Figure 3.** Results of a bootstrapping analysis performed over all reactivation-based analyses reported in the manuscript. To ensure that our reactivation measures and analyses were robust, we performed a bootstrapping analysis in which we sampled with replacement from our data (n=39), and repeated each analysis 1000 times. Specifically, on each re-sampled dataset, we fit the same set of mixed effects models used on the full sample, which yielded a nonparametric estimate of the sampling distribution for each effect. All effects replicated, and the effect of Fan Out condition reactivation scores on transfer choice reaction times strengthened with this analysis. Black dotted lines represent where 95% density of these distributions lies, and red dotted lines are drawn at 0. 95% intervals that exclude zero are considered significant.
